# GlioVision: A Multi-Modal MRI Framework for Non-Invasive Glioma Molecular Biomarkers Prediction

**DOI:** 10.64898/2026.04.22.719993

**Authors:** Anam Nazir, Muhammad Nadeem Cheema, Yu-Chun Hsu, Xiaoqian Jiang, Jay-Jiguang Zhu, Akdes Serin Harmanci, Arif Harmanci

## Abstract

Gliomas are aggressive primary brain tumors that necessitate critical molecular biomarker predictions for optimal clinical decision-making. Traditional assessment relies on surgical tumor specimens analysis, which carries procedural risks and sampling bias due to tumor heterogeneity. Existing deep learning methods for non-invasive prediction lack real-time applicability, remain resource-intensive, and are frequently trained on narrowly represented datasets. We present GlioVision, a framework built on the MONAI library to process multimodal data, including glioma MRI and molecular labels, to predict and identify, non-invasively, four major glioma molecular biomarkers: IDH mutation, 1p/19q co-deletion, MGMT methylation, and WHO grade. The core architecture comprises Spatially and Channel-wise Recalibrated 3D DenseNet (SCRU-DenseNet), which utilizes a computational attention gate and an Adaptive Contrast-Specific Processing Stream (ACPS) to tackle multi-site, heterogeneous datasets. We introduced the Confidence-Filtered Predictive Manifold (CFPM) to manage uncertainty by excluding predictions with low confidence. GlioVision is trained and validated on the largest multi-cohort datasets, achieving strong biomarker prediction with AUCs of (IDH 0.94, 1p/19q 0.87, MGMT 0.86, WHO grades 0.92), supporting molecularly defined glioma diagnosis under the WHO 2021 classification guidelines. Finally, we provide a Differential Training Integrity Assessment (DTI-A) to analyze routes of MRI data privacy protections through model obfuscation. Our results advance the codebase, model release, and leakage considerations around MRI data analysis literature.

## 1. Introduction

Gliomas are the most common primary parenchymal brain tumors, including astrocytoma, oligodendroglioma, and ependymoma [1]. Astrocytoma consists of grades 1, 2, 3, and 4 tumors. Glioblastoma (GBM) is the highest-grade tumor among astrocytoma with wild-type isocitrate dehydrogenase (IDH), with a five-year survival rate of only 6.9% [2]. Patient outcomes depend on pathological diagnoses, which are heavily reliant on molecular changes in DNA, RNA, proteins, and epigenetics, tumor location, functional status, age, and therapies received. Currently, tumor tissue analysis by pathologists, aided by molecular technologies such as methylation profiling and next-generation sequencing, is the only way to identify the specific molecular mutations (such as IDH mutation, 1p/19q co-deletion, MGMT methylation) necessary for pathological diagnosis.

Obtaining tumor tissue either by craniotomy or by tumor biopsy is invasive and poses risks. Furthermore, molecular testing of tumor tissue is time-consuming (Turnaround time is about 10 to 21 days) and resource intensive as well as presenting sampling bias give the heterogeneity of GBM. Magnetic Resonance Imaging (MRI) [3], including diagnosis, surgical planning, and response assessment. Artificial intelligence with deep learning (DL) methods has demonstrated promising potential for non-invasive characterization of glioma biomarkers from MRI scans [4].

While preliminary models can predict multiple biomarkers, many remain computationally intensive, struggle to account for the full range of tumor categories, molecular markers, but often struggle to generalize due to small or homogeneous training datasets [5]. These limitations hinder clinical adoption, as models must deliver high accuracy with low computational demands to be practical in real-world workflows. Additionally, numerous preliminary architectures claim to offer advanced solutions for glioma biomarker prediction; however, they have limited practicality and reproducibility [5], [6], [7], as they often lack practical testing scenarios or the means to verify the authenticity of the claimed methods [8].

Modern MRI imaging provides high-resolution spatial information about tumor morphology, including shape, size, and anatomical location [9]. Key molecular biomarker labels from TCIA datasets, IDH mutation status, deletion of the short arm of chromosome 1 and the long arm of chromosome 19 (1p/19q codeletion), O^6^-Methylguanine-DNA Methyltransferase (MGMT) promoter methylation, and tumor grade, serve as an ordinal indicator of malignancy severity (e.g., grade 2–4) and are typically encoded as binary variables indicating mutation presence, chromosomal alteration, or epigenetic modification. These labels serve as complementary categorical and ordinal data for a deep learning model. The World Health Organization (WHO) 2021 classification further formalized this approach by defining glioma entities primarily based on molecular features, particularly IDH mutation status, 1p/19q codeletion, and tumor grade [2]. By integrating multimodal input imaging and clinical data into a unified predictive framework, we have proposed GlioVision, which enables improved prediction of glioma biomarkers. GlioVision processes four types of 3D, multi-sequence MRI scans: FLAIR, T1, T1-GD, and T2, combined with clinical molecular labels.

GlioVision’s core architecture is based on spatially and channel-wise recalibrated three-dimensional (3D) Dense Convolutional Network, termed SCRU-DenseNet. This is a modified 3D DenseNet with a Squeeze-and-Excitation (SE) module, designed to learn vital imaging patterns while maintaining computational feasibility for real-time use. At the core of the system, the proposed SCRU-DenseNet introduces a feature-importance weighting strategy, where the SE module serves as a computational attention gate, dynamically modeling channel-wise interdependencies through global average pooling and a sigmoid-gated reduction–expansion cycle. To further enrich the network’s representational capacity, we integrate an Adaptive Contrast-Specific Processing Stream (ACPS) that treats each MRI sequence and tumor mask as independent input streams, enabling modality-specific training and substantially broadening the input feature space. This design strengthens the extraction of radiogenomics signatures by capturing both deep intra-tumoral and broad peri-tumoral contextual information, thereby improving generalizability across institutions.

To ensure fair prediction across all biomarkers and overcome the lack of available DenseNet weights, we trained GlioVision from scratch on the largest multi-cohort glioma dataset (over 2279 subjects), utilizing a dynamic weighted loss strategy with staged learning to balance underrepresented classes and facilitate efficient real-time deployment on standard GPUs. Traditional biomarker prediction models generate outputs for all cases regardless of confidence, which may lead to clinically unsafe misclassifications. GlioVision proves itself a fail-safe model by integrating a Confidence-Filtered Predictive Manifold (CFPM) to clinical risk [10] that selectively withholds low-confidence predictions. GlioVision also introduced a Differential Training Integrity Assessment (DTI-A), a privacy-preserving model obfuscation strategy, to minimize the risk of unauthorized data misuse when sharing. The following are the key contributions of the proposed model:

1. We present GlioVision, a MONAI-based framework designed to process multimodal data, including glioma MRI and molecular labels, for non-invasive prediction of four key glioma molecular biomarkers: IDH mutation, 1p/19q co-deletion, MGMT methylation, and WHO grade.
2. We introduce a modality-specific training strategy where each MRI sequence and tumor mask is treated as an independent input stream. Using the Adaptive Contrast-Specific Processing Stream (ACPS), the model learns tumor-core and surrounding-tissue features, improving GlioVision’s predictive performance.
3. The core architecture SCRU-DenseNet achieves feature importance weighting based on the SE module, which functions as a computational attention gate that dynamically computes the channel-wise interdependence of latent feature maps by performing global average pooling followed by a sigmoid-gated dimensionality reduction/expansion cycle.
4. Our confidence-based CFPM module is an adaptive post-processing unit filtering strategy that accepts only high-certainty predictions and systematically omits unreliable outputs to improve reliability.
5. Our privacy-preserving strategy (DTI-A) addresses privacy concerns using induced controlled Gaussian noise directly within the multi-parametric MRI for model obfuscation. This process allows us to assess the re-identification of the patient data and the intellectual property of the learned radiogenomics feature space.

Predicting IDH mutation status, 1p/19q co-deletion, and MGMT promoter methylation pre-operatively informs the immediate surgical plan and post-surgical treatment e.g., choice of chemotherapy and radiation targets). GlioVision’s ability to provide this pathological prediction without an initial biopsy is a major clinical gain.

## 2. Methods

This section describes the methodological framework developed for this study. Beginning with the design of the SCRU-DenseNet architecture, including its channel-recalibration mechanism, Adaptive Contrast-Specific Processing Stream (ACPS), followed by a description of the multi-institutional dataset assembled for model development and evaluation.

GlioVision is a multi-modal architecture that combines heterogeneous data sources from the Cancer Imaging Archive (TCIA) database [11] to integrate multi-sequence MRI scans (T1, T1GD, T2, FLAIR). The imaging component captures detailed spatial information on tumor morphology, including size, shape, and anatomical location, while the non-imaging component incorporates structured clinical and genomic features, covering four key biomarkers. To standardize input data, an organized multi-step pipeline was implemented, shown in Figure 1. Raw DICOM scans were converted to NIfTI format using dcm2niix [12] and registered to the MNI152 standard space with Elastix (affine + nonrigid) [13]. Intensity nonuniformities were corrected using N4 bias correction, and HD-BET was applied for brain extraction and skull stripping [14].

**Figure 1:**
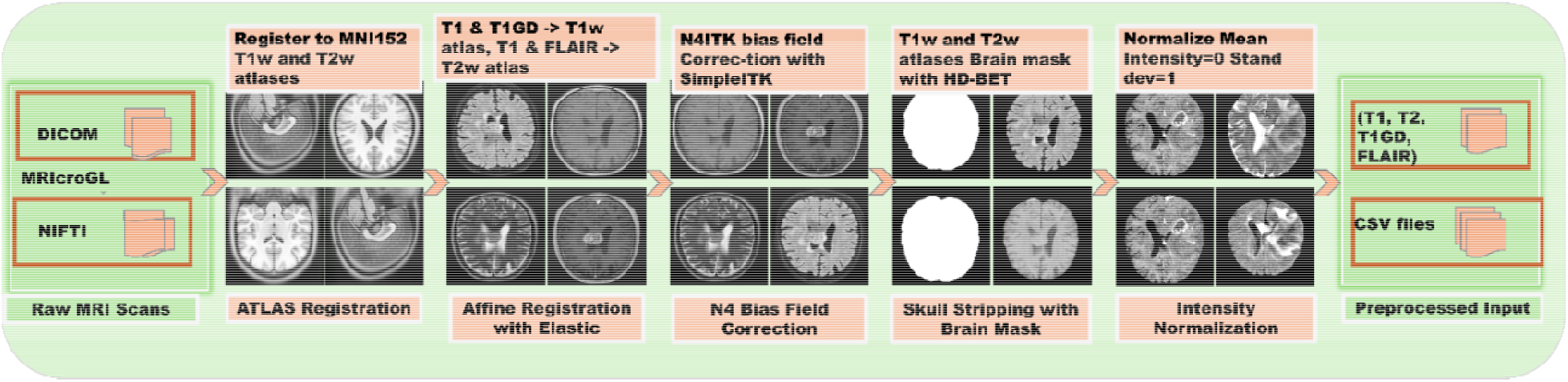
Preprocessing pipeline

### 2.1. Adaptive Feature Synthesis via Multi-Sequence Contextual Augmentation

We introduce adaptive feature synthesis via multi-sequence contextual augmentation, a critical component of the adaptive contrast-specific processing stream (ACPS), as shown in Figure 2, designed to significantly enrich the input feature space for improved molecular stratification. This method operates through modality-specific training [15], [16]treating each MRI sequence as a separate, independent input stream, this segregation ensures that the unique tissue contrast mechanisms and pathology-sensitive features inherent to each sequence are fully utilized and disentangled from one another [15].

**Figure 2:**
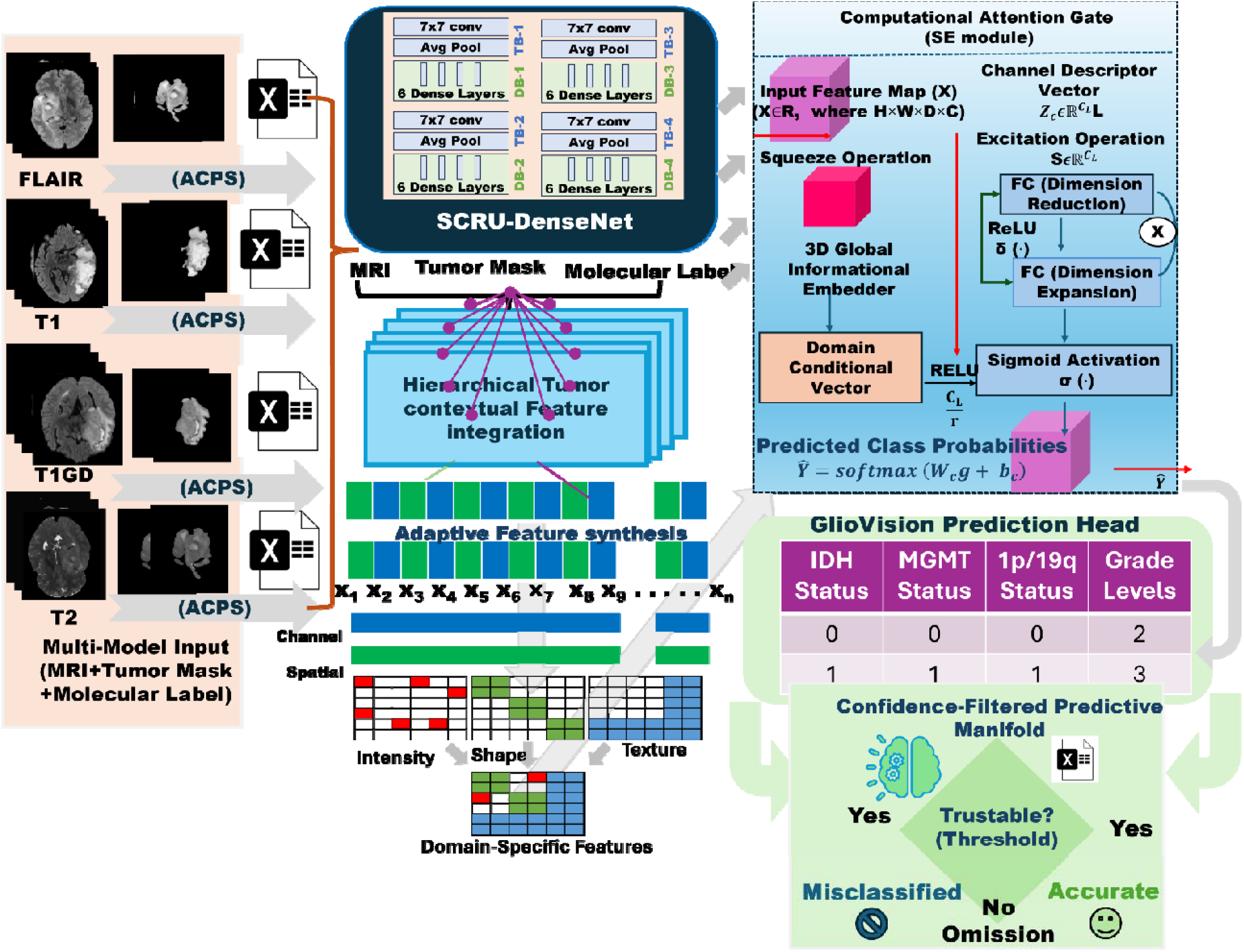
The GlioVision pipeline begins with the processes of multi-parametric, multi-modal input MRI, clinical label, and tumor masks, utilizing multi-modal training with the Adaptive Contrast-Specific Processing Stream (ACPS), which focuses feature extraction on the relevant core volume. Features are then routed to the Spatially and Channel-wise Recalibrated 3D DenseNet (SCRU-DenseNet), which uses a Computational Attention Gate to dynamically recalibrate channels. These processed feature embeddings are fused via Hierarchical Tumor-Contextual Feature Integration (HTC-FI), which integrates both local tumor characteristics and global contextual information into a single, unified representation. Finally, the GlioVision prediction head generates biomarker predictions, which are then passed to the Confidence-Filtered Predictive Manifold (CFPM) to minimize clinical misclassification risk by excluding low-confidence prediction results.

To achieve superior feature granularity, we integrate the tumor segmentation mask as a contextual augmentation channel accompanying each modality input. This creates a Hierarchical Tumor-Contextual Feature Integration (HTC-FI) mechanism that simultaneously captures high-resolution features, explicitly focused on the lesion’s microenvironment and heterogeneity, informed by the mask and features describing the functional relationship and mass effect of the tumor relative to the whole brain volume. For further details, see Supplementary document (section S1).

Each MRI sequence provides unique insights into tissue contrasts and pathological details individually. This technique exploits the diverse, complementary information intrinsic to multi-parametric MRI to improve generalization performance in the presence of tumor variability and scanner differences. Furthermore, it reduces the dependence on explicit tumor segmentation by enabling the model to extract whole-volume contextual representations informed by modality-specific contrast. To optimize computational resources and focus on the most relevant brain regions, the outermost 15 slices at each end of every 3D volume were excluded from the training process. The integration of ACPS with the SCRU-DenseNet backbone follows a late-fusion approach. Specifically, each MRI sequence and its corresponding tumor mask are concatenated to form a dual-channel input for modality-specific encoders. This segregation allows the model to disentangle pathology-sensitive features unique to each contrast mechanism (e.g., edema in FLAIR vs. necrosis in T1GD). The resulting independent feature maps are then unified via a channel-wise concatenation layer.

### 2.2. Spatially and Channel-wise Recalibrated 3D DenseNet (SCRU-DenseNet) Unit

The core feature extraction engine of GlioVision is the SCRU-DenseNet as demonstrated in Figure 2, a specialized architecture engineered for the efficient modeling of volumetric, multi-institutional MRI data. This design represents a significant methodological advancement over conventional 3D Convolutional Neural Networks (3D CNNs), addressing the inherent computational burden and feature redundancy associated with large-scale medical imaging. We adopt a modified SCRU-DenseNet represented in Figure 2, which serves as the feature extractor for volumetric information from multi-sequence brain MRI data *X*∈*R*, where *H×W×D×C* defines the isomorphous tensor space, and *C* denotes the cardinality of the multi-parametric image acquisition set. The SCRU-DenseNet is fundamentally rooted in a dense connectivity paradigm, where each layer within a dense block receives feed-forward input from all preceding layers and contributes its feature maps to all subsequent layers. The SCRU-DenseNet comprises L densely connected blocks, with each block creating feature maps that are fused and fed through nonlinear transformations to generate the final feature tensor *F(L)*∈*R*, where *H*_*L*_*w*_*L*_*D*_*L*_denote the spatial dimensions after reduction by pooling and convolutional strides, with *c*_*L*_ denoting the output channel count.

### 2.3. The Computational Attention Gate

The primary contribution lies in integrating a computational attention gate, implemented via a modified 3D Squeeze-and-Excitation (SE) module, into the output of each dense block. This unit performs spatio-channel recalibration to dynamically adjust the importance of feature maps before they are fed into the next layer.

1. Squeeze Operation: Figure 2 reveals that global information embedding is performed using the squeeze operation. This step effectively aggregates global spatial information into a local descriptor. This mechanism conducts channel-wise recalibration conditioned on a learned domain feature vector *d*_*C*_*∈R*^*K*^ corresponding to cohort *i*, where *K* is the embedding dimensionality. Recalibration is initiated by reducing the spatial dimensions of 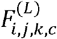 through global average pooling, producing a channel descriptor vector 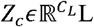, where each component *z*_c_ is derived as the average value of features across all spatial positions

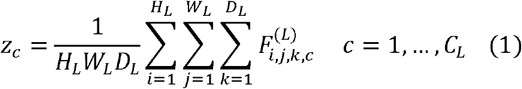
2. Excitation operation is done through channel-wise interdependence. The excitation stage assimilates domain information by projecting *z* and *d*_*i*_ linearly into a lower-dimensional embedding space of size 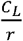, with r acting as the reduction ratio to control model capacity and regularization. Formally, the excitation vector 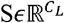 is acquired through a nonlinear transformation activated by ReLU δ(⋅) followed by a sigmoid activation σ (⋅) to the sum of the projections:

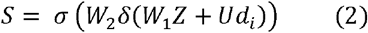

Where 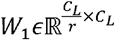 and 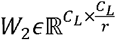 are the learned weights of the two fully connected layers, and 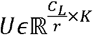 projects the domain embedding into the reduced channel space. The recalibration vector is then propagated over the spatial dimensions and applied as an element-wise multiplier to the original feature tensor, resulting in the recalibrated feature map.

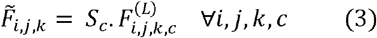

Thereby shaping channel activations informed by both learned feature weights and domain factors. Finally, the recalibrated features are spatially aggregated via global average pooling into a fixed-length vector 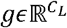, which is input to a fully connected classification head with a weight matrix 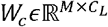 and bias 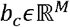, where M is the number of biomarker classes. The model outputs predicted class probabilities ŷ through a softmax function:

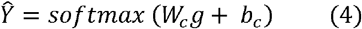

Given the natural imbalance in the clinical labels, an adaptive class-weighted loss function was used. This approach imposes higher penalties for misclassifications of rare classes and adjusted class weights at each training epoch to track shifts in the label distribution, supporting balanced model training. Since no pre-trained weights existed for volumetric DenseNet-121 in this domain, the model was trained from scratch. To accommodate the GPU memory constraint (8 GB), we adopted a multi-stage hyperparameter tuning approach coupled with an adaptive learning rate schedule. The final prediction is a probabilistic vector across biomarker classes.

### 2.4. Data Collection and Cohort Assembly

A custom-assembled, multi-cohort dataset comprising over 2279 unique subjects was assembled for our study evaluation, as represented in

Table *1*. Data collection involves several publicly accessible databases from the TCIA sites, including UCSF-PDGM [17], UPENN-GBM [18], TCGA-LGG [11], TCGA-GBM [19], EGD [20], IvyGAP [21], and RHUH-GBM [22]. The dataset includes 2,279 subjects, contributing a total of 9,116 MRI scans and approximately 1,412,980 image slices. For more details, see Supplementary document (section S4). These data were split into training, validation, and test sets at 70/15/15 for each biomarker across subjects. The 70/15/15 partitioning was executed at the subject level using unique patient identifiers to ensure that no image data (slices or volumes) from a single subject appeared in more than one set. This approach maintains the statistical integrity of the 70/15/15 ratio within each predictive task while ensuring that the test cohorts consist entirely of previously unseen subjects, thereby eliminating the risk of data leakage.

**Table 1:** Dataset with biomarker distribution details for IDH and 1p/19q, MGMT methylation, and WHO grades for EGD, TCGA LGG and GBM, UCSF-PDGM, UPENN-GBM, RHUH-GBM, and IvyGAP.

| Data Cohorts | Subjects | IDH (0) | IDH (1) | MGMT (0) | MGMT (1) | 1p/19q (0) | 1p/19q (1) | Grade 4 | Grade 3 | Grade 2 |
| --- | --- | --- | --- | --- | --- | --- | --- | --- | --- | --- |
| EGD | 774 | 312 | 155 | 0 | 0 | 186 | 73 | 502 | 79 | 135 |
| TCGA LGG<br>GBM | 164 | 90 | 56 | 0 | 0 | 147 | 12 | 100 | 37 | 27 |
| UCSF-PDGM | 491 | 390 | 101 | 296 | 112 | 385 | 15 | 394 | 42 | 55 |
| UPENN-GBM | 671 | 546 | 125 | 121 | 170 | 0 | 0 | 0 | 0 | 0 |
| RHUH-GBM | 40 | 36 | 4 | 0 | 0 | 0 | 0 | 40 | 0 | 0 |
| IvyGAP | 39 | 31 | 6 | 0 | 0 | 27 | 3 | 36 | 2 | 0 |
| TOTAL | 2279 | 1405 | 447 | 417 | 282 | 745 | 103 | 1103 | 180 | 254 |

## 3. Results

The model was trained for 300 epochs using the Adam optimizer with stable optimization (learning rates ranging from 1e-6 to 5e-6, weight decay from 1x 10^-6^ to 1x 10^-4^, and batch sizes of 10). All experiments were conducted on a system equipped with an NVIDIA RTX 4060 GPU (8 GB of VRAM), an Intel Core i5-12th-generation processor, 16 GB of DDR4 RAM, and an Ubuntu 22.04 LTS (64-bit) operating system. Training times varied based on the selected model’s biomarker, dataset size, and number of epochs, ranging from 24 to 36 hours for more demanding tasks.

As shown in Figure 3 and Table 2, our model achieves an AUC of 0.94 for IDH, with higher precision and recall for wild-type samples, indicating areas for improvement in the identification of mutant samples. In the 1p/19q codeletion task, as represented in Figure 3, the model achieved an AUC of 0.86, with Class 1 exhibiting high sensitivity. For MGMT promoter methylation, performance was balanced and clinically relevant with an AUC of 0.86. Across tumor grades 2, 3, and 4 from Figure 3, AUCs were 0.92, 0.94, and 0.94, respectively.

**Table 2:** Results for GlioVision in terms of precision, recall, sensitivity, and specificity for the four major glioma key molecular biomarkers.

| Metric | IDH<br>Class 0 | IDH<br>Class 1 | 1p/19q<br>Class 0 | 1p/19q<br>Class 1 | MGMT<br>Class 0 | MGMT<br>Class 1 | Grade<br>Class 2 | Grade<br>Class 3 | Grade<br>Class 4 |
| --- | --- | --- | --- | --- | --- | --- | --- | --- | --- |
| Precision | 0.88 | 0.8 | 0.967 | 0.5 | 0.8451 | 0.8943 | 0.7024 | 0.7097 | 0.7097 |
| Recall | 0.93 | 0.7 | 0.8286 | 0.8571 | 0.8219 | 0.9091 | 0.8182 | 0.6875 | 0.7132 |
| F1 Score | 0.91 | 0.75 | 0.8923 | 0.6316 | 0.8333 | 0.9016 | 0.7559 | 0.6984 | 0.711 |
| Sensitivity | 0.9444 | 0.7843 | 0.8286 | 0.8571 | 0.8219 | 0.9091 | 0.8182 | 0.6875 | 0.7132 |
| Specificity | 0.7843 | 0.9444 | 0.8571 | 0.8286 | 0.8451 | 0.8943 | 0.9584 | 0.9556 | 0.9556 |

**Figure 3:**
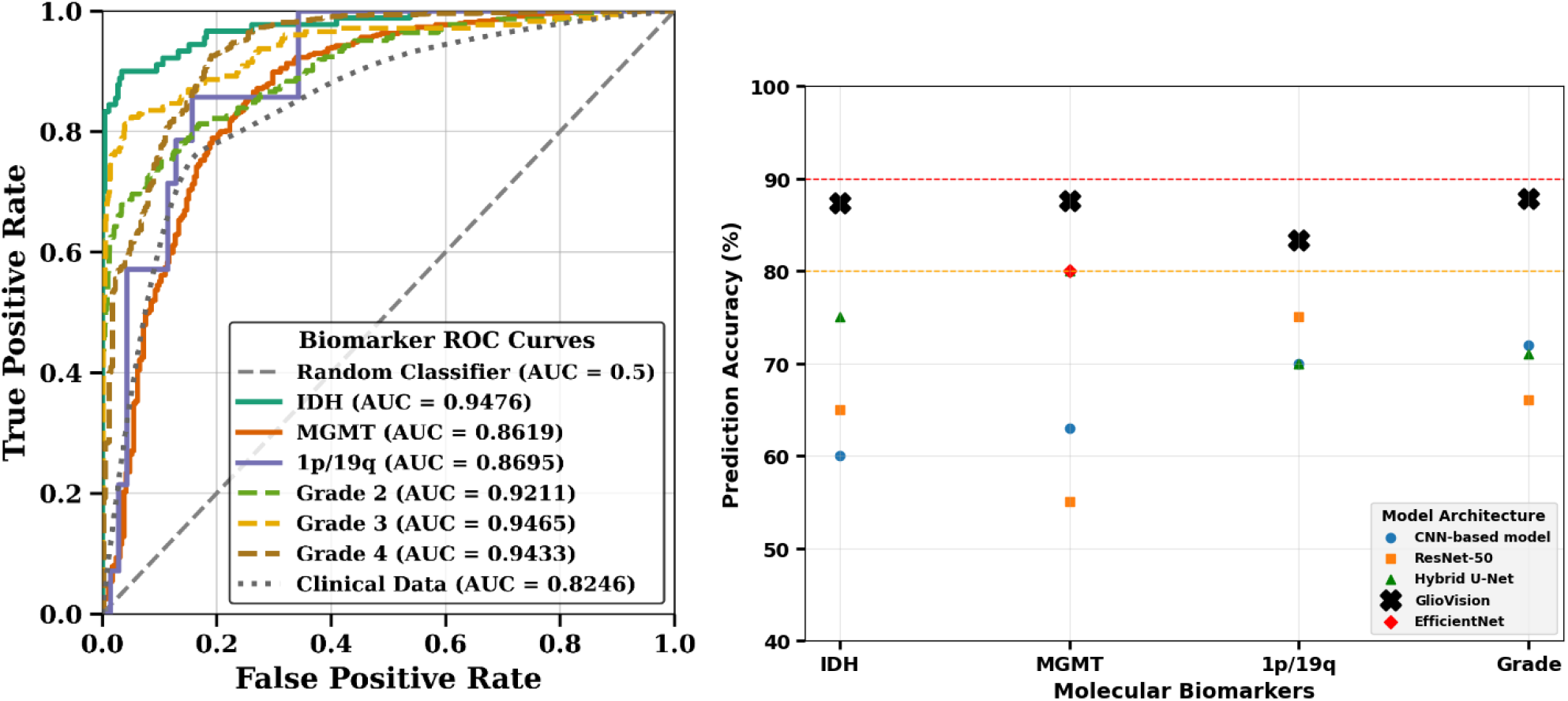
Left: AUC curves for four biomarkers on public and clinical datasets. Right: Comparison of GlioVision with state-of-the-art methods using the same training, testing, and validation cohorts under identical settings.

The efficacy of GlioVision was determined by quantifying the difference, as shown in

Table 3 between the model accuracy (A model, which represents the observed rate of correct predictions), and the baseline accuracy (*A* _*baseline*_, which represents the null hypothesis defined as the accuracy achieved by merely predicting the majority class *(Size of the Majority Class/Total Samples)*). The magnitude of this difference serves as the true measure of generalizable feature learning. The *P-value* uses a *Z-test* to compare *(A* _*model*_) against *(A* _*baseline*_*) and determines* whether the performance is statistically superior to guessing. A result of *P < 0*.*001* confirms that our model achieved its accuracy by genuinely learning features, not random chance

**Table 3:**
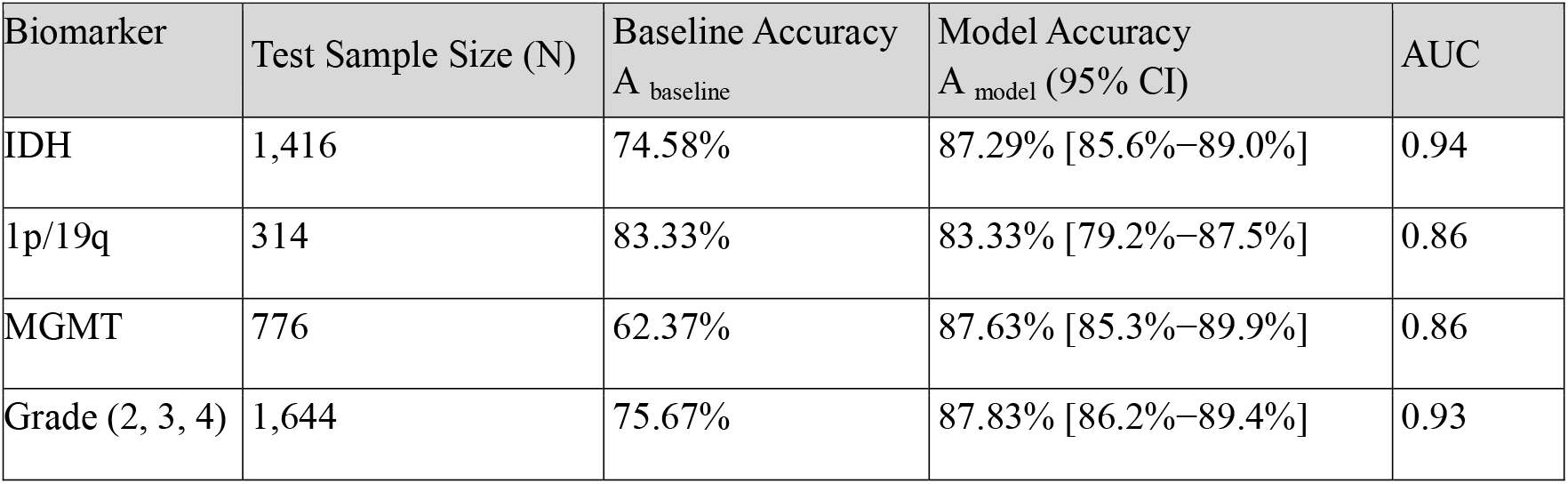
Performance of GlioVision across different biomarkers. The table displays test sample sizes, baseline accuracy, model accuracy with 95% confidence intervals, statistical significance (p-values), and AUC values for IDH, 1p/19q, MGMT, and tumor grades (2, 3, 4).

| Biomarker | Test Sample Size (N) | Baseline Accuracy<br>$A_{baseline}$ | Model Accuracy<br>$A_{model}$ (95% CI) | AUC |
| --- | --- | --- | --- | --- |
| IDH | 1,416 | 74.58% | 87.29% [85.6%–89.0%] | 0.94 |
| 1p/19q | 314 | 83.33% | 83.33% [79.2%–87.5%] | 0.86 |
| MGMT | 776 | 62.37% | 87.63% [85.3%–89.9%] | 0.86 |
| Grade (2, 3, 4) | 1,644 | 75.67% | 87.83% [86.2%–89.4%] | 0.93 |

GlioVision successfully classified four major glioma biomarkers, with the IDH biomarker demonstrating the highest performance, with an Area Under the Curve (AUC) of 0.9476. To validate the reliability of these estimates, 95% *Confidence Intervals (CIs)* were calculated to quantify precision as a function of the test sample size *N*. For example, the grade model’s accuracy was tightly bound between 86.2% and 89.4%. Furthermore, the significance of the results was confirmed by comparing the *P-value* against the baseline of the majority class. All four glioma biomarkers, including MGMT, which achieved the greatest practical gain by increasing performance from a low baseline of 62.37% to 87.63%, demonstrated statistically significant improvement *(P < 0.001)*.

Conversely, the 1P/19q biomarker, despite achieving strong technical discrimination (AUC = 0.87), shows statistical significance against its high 83.3% majority-class baseline, with a class 1 sensitivity of 85.7%. This result is attributed to the strict statistical test and the small sample size of 314. In contrast, GlioVision demonstrates strong discriminative capacity, with an AUC of 0.8695, indicating that the model has successfully learned the complex radiomic signatures of 1p/19q status rather than relying on class frequency. This is further evidenced by a balanced sensitivity of 85.71%, which ensures that nearly 9 out of 10 oligodendroglioma patients are correctly identified. To quantify the real-time applicability of the GlioVision framework, we benchmarked inference speed on a standard 8-GB GPU workstation. The model demonstrated an average inference time of 0.040 seconds (40 ms) per patient study.

To assess the real-world applicability of the GlioVision model, we further evaluated its performance on an independent, clinically acquired cohort from an affiliated hospital (Houston, Texas, USA). This validation set comprised 57 patients with matched MRI scans and confirmed IDH status, as determined from variant calling files (VCFs). The IDH mutation status was determined by examining the presence of the two clinically relevant hotspots: chr2:209113112 (R132H) and chr2:209113113 (R132C). As shown in the AUC curve in Figure 3. The model achieved an AUC of 0.8246 for IDH.

### 3.1. Diagnostic Alignment with WHO 2021 Guidelines

It is important to note that the primary objective of this study is not direct tumor subtyping, but rather the prediction of key molecular biomarkers, particularly IDH mutation status, 1p/19q codeletion, MGMT methylation, and tumor grade. These biomarkers form the basic components required for glioma classification under the current WHO 2021 guidelines. Therefore, even though tumor subtype definitions have changed over time, our model remains applicable because it predicts the fundamental molecular features used in modern diagnosis. To achieve clinical alignment of GlioVision predictions, we follow the guidelines aligned with the WHO 2021 (5th Edition) classification of tumors of the Central Nervous System [23]. This framework notes that molecular markers, particularly IDH mutation status, replace histological grading in determining the final integrated diagnosis.

A total of 64 subjects from the test cohort were analyzed, and both predicted IDH status and histological grade are shown in Table 4. The predictions were mapped to the WHO-2021 diagnostic. For IDH wildtype cases. Among 41 subjects predicted to be IDH-wildtype, the model classified 36 (87.8%) as Grade 4, corresponding directly to the WHO-2021 category Glioblastoma, IDH-wildtype, and 5 subjects (12.2%) as Grade 2 or Grade 3. Under the WHO-2021 molecular-first guidelines, these 5 cases are reclassified as molecular glioblastomas, with an IDH-wildtype molecular profile indicating high-grade biological behavior despite lower histological grade predictions. In this study, the term molecular glioblastoma refers to tumors predicted to be IDH-wildtype and histologically Grade 2 or 3. Under the WHO 2021 criteria, these tumors may be reclassified as Glioblastoma, IDH-wildtype if additional defining molecular alterations are present; otherwise, they may be categorized as Not Elsewhere Classified (NEC). For the IDH-mutant cases, of the 23 subjects predicted to be IDH-mutant, all fell within the WHO-2021 framework for Astrocytoma, IDH-mutant (Grades 2–4). Notably, 9 subjects were classified as grade 4, consistent with the newly recognized category Astrocytoma, IDH-mutant, Grade 4, which replaces the obsolete IDH-mutant glioblastoma designation from earlier classifications.

**Table 4:** Diagnostic results for the Grade aligned with the WHO guideline for the classification of glioma.

| Predicted IDH Status | Predicted Grade | WHO-2021 Category | No of Subjects |
| --- | --- | --- | --- |
| Wildtype | Grade 4 | Glioblastoma, IDH-wildtype | 36 |
| Wildtype | Grade 2/3 | Molecular Glioblastoma | 5 |
| Mutant | Grade 2/3/4 | Astrocytoma, IDH-mutant | 23 |

For a subset of 29 subjects for whom predictions of IDH status, 1p/19q codeletion, and histological grade were available, the diagnostic analysis is shown in Table 5. The model identified 15 subjects as Glioblastoma, IDH-wildtype (Grade 4), 3 subjects as Molecular Glioblastoma (IDH-wildtype, Grade 2/3), 7 subjects as Oligodendroglioma, IDH-mutant and 1p/19q-codeleted, 2 subjects as Astrocytoma, IDH-mutant (non-codeleted), and 2 subjects as atypical or biological outliers. Among the 19 subjects classified as Glioblastoma, IDH-wildtype (including molecular glioblastomas), 15 cases (78.9%) were predicted to have MGMT promoter methylation, a key biomarker associated with improved response to temozolomide therapy.

**Table 5:** Integrated WHO-2021 Diagnosis for IDH, Grade, and 1p/19q.

| Predicted IDH Status | Predicted 1p/19q | Predicted Grade | WHO-2021 Integrated Diagnosis | No of Subjects |
| --- | --- | --- | --- | --- |
| Wildtype | - | 4 | Glioblastoma, IDH-wildtype | 15 |
| Wildtype | - | 2/3 | Molecular Glioblastoma | 3 |
| Mutant | Codeleted | 2/3 | Oligodendroglioma, IDH-mutant | 7 |
| Mutant | Non-Codeleted | 2/3/4 | Astrocytoma, IDH-mutant | 2 |

### 3.2. Comparison with State-of-the-Art Methods

There are several glioma biomarker prediction models available, but many of the published techniques either do not provide the source code for reproduction [8] or the provided code relies on outdated libraries and requirements of computationally infeasible GPU resources [24], making them difficult to reproduce or insufficient for training and testing on large cohorts. However, we still tested our customized data cohort against available models, using the same settings we use for GlioVision. The GlioVision shows improvements over conventional architectures, as shown in Figure 3, across trained models, including CNN-based model [25], ResNet-50 [7], a hybrid U-Net [24], a radiomics-based method [6], and EfficientNet [26]. For example, GlioVision reaches 87.0% accuracy for IDH prediction, surpassing CNN-based model [25] 60.0% and the hybrid U-Net’s [24] 75.0%. Similarly, for 1p/19q codeletion, it achieves 83% accuracy, and for MGMT methylation, GlioVision’s 86.19% accuracy exceeds that of the best baseline, EfficientNet, at 79.6%. For glioma grading, GlioVision’s results are notable, with 87.87% accuracy, outperforming the 65% accuracy achieved by hybrid U-Net models under our settings and configurations.

### 3.3. Addressing Misclassification with Confidence-Filtered Predictive Manifold (CFPM)

Typically, biomarker prediction frameworks provide a prediction every time, despite having low prediction certainty, which may result in misguided clinical decisions. GlioVision proves itself a fail-safe model by integrating a CFPM that selectively withholds low-confidence predictions. Within this framework, the model outputs probabilities for each class, and the maximum predicted probability (*Max Prob*) is extracted as a trust score. If *Max Prob* exceeds a predefined confidence threshold (e.g., 0.8, depending on the clinical utility), the predicted class is accepted. Otherwise, the prediction is omitted to prevent low-confidence outputs from being acted upon. Formally, we can define CFPM according to our scenario with a probability vector as:

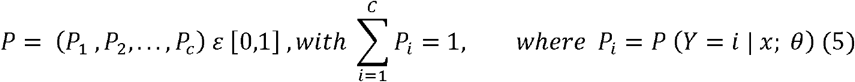

P is the predicted posterior probability of class *i* given input *x* and model parameters *θ*. Here we can define the maximum predicted probability (trust score *s*) as:

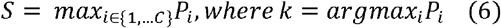

Where the indicator factor 𝕝_{*f*}_ is defined as:

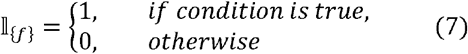

Given a confidence threshold τ *∈*[0,1], define the acceptance function:

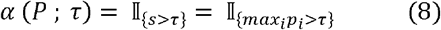

Then, the final decision function is:

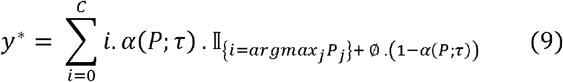

Equivalently, this can be written as:

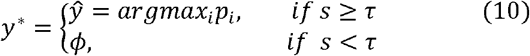

where denotes omission of the prediction to prevent low-confidence output. Additionally, this defines a thresholder, a nonlinear operator C→{1,,C}∪{ } that acts as a self-regulating filter:

In our experimental evaluation, shown in Figure 4 (a, e) and Supplementary documentation (section S6), applying CFPM over GlioVision for IDH prediction led to 15 out of 92 predictions being omitted, including 10 originally incorrect and 5 correct but uncertain cases. This reduced the total number of accepted predictions to 77 but improved the accuracy on accepted samples from 85.87% to 92.21%, while cutting misclassifications by more than half (from 13 to 6).

**Figure 4:**
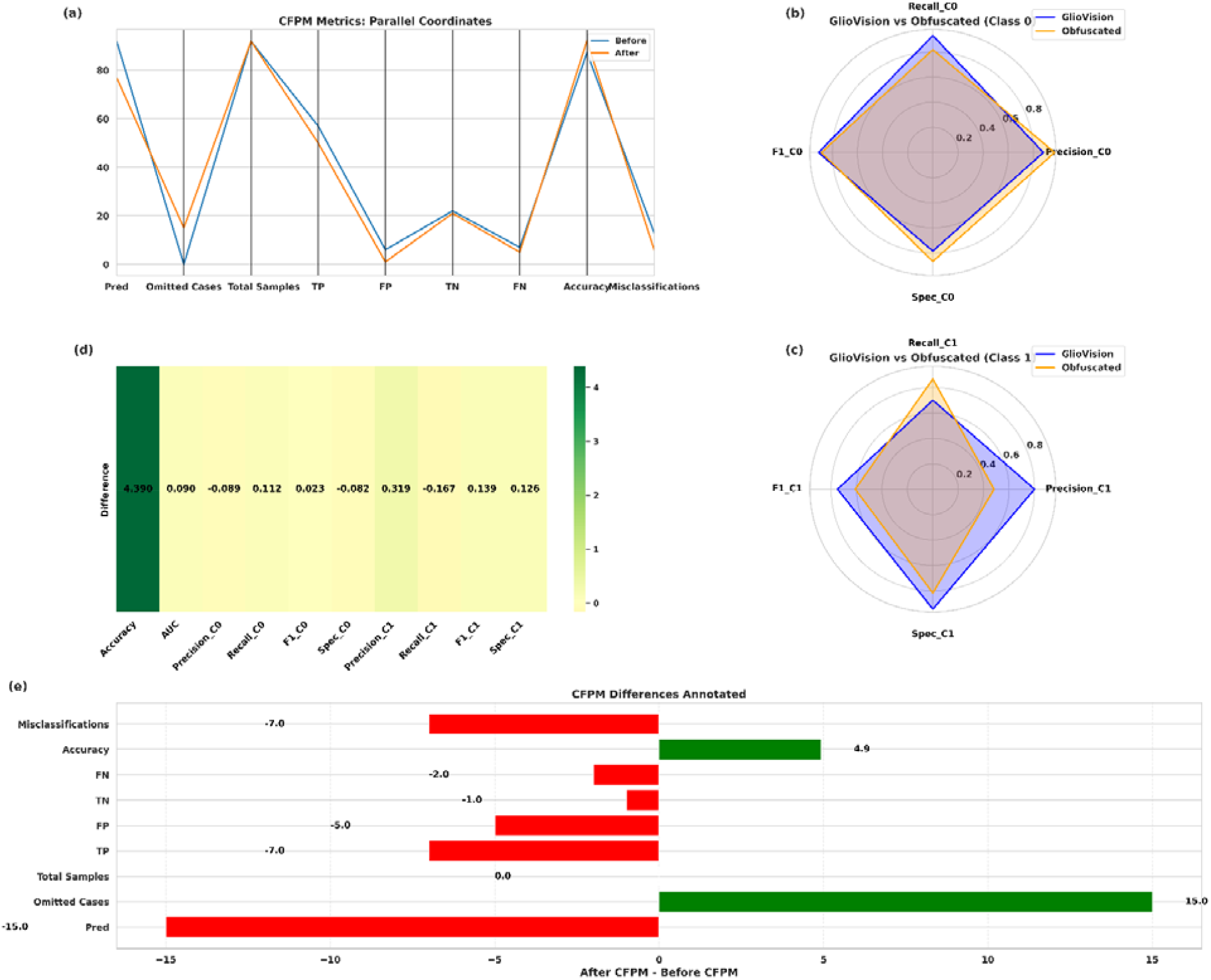
(a) Parallel coordinates plot of CFPM metrics showing the shift from before to after CFPM, including predictions used, omitted cases, total samples, true positives, false positives, true negatives, false negatives, accuracy, and misclassifications. (b, c) Radar plots compare GlioVision versus Obfuscated performance for Classes 0 and 1, displaying precision, recall, F1-score, and specificity, with lower axis labels for readability. (d) A heatmap of GlioVision differences highlights positive improvements in green and declines in red, while (e) a horizontal bar chart illustrates CFPM metric differences, showing improvements (green) and declines (red) across all metrics.

### 3.4. MRI Privacy Assessment in GlioVision via Model Obfuscation with (DTI-A)

To assess the feasibility of providing privacy protections in MRI data analysis [27], we have examined controlled data perturbations intended to mask sensitive training data while preserving clinically acceptable model performance [28]. Our strategy, DTI-A, directly reshapes the model’s learned patterns by retraining on perturbed samples, thereby reducing the risk of proprietary or patient-identifiable image data being inferred from the model’s weights. First, the baseline model, as represented in Figure 5 is trained on the original MRI input dataset to learn representative features for prediction tasks. After the initial model has been trained, its weights serve as the foundation for obfuscation. Subsequently, a perturbed version of the dataset is generated by adding carefully controlled Gaussian noise exclusively to brain tissue regions, without altering the background. For instance, each axial slice *z* of the MRI volume *I* and a brain mask *M*_z_ is created by thresholding the intensity values:

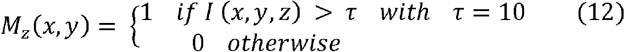

where *I* (*X,y,z*)is the original voxel intensity at spatial coordinates (*x, y*) in slice *z*. τ is the hreshold value (10). *M*_*z*_(*x, y*) is 1 where the voxel intensity is above the threshold (inside brain tissue) and 0 otherwise (background). Next, Gaussian noise *N*_*z*_ with zero mean and standard deviation proportional to the slice’s own standard deviation is generated and added only within the brain mask:

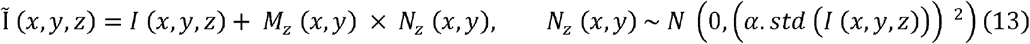

where Ĩ (*x,y,z*) is the new, perturbed voxel value. *N*_*z*_ (*x, y*) is random noise sampled from a Gaussian (normal) distribution with mean 0 and standard deviation equal to *α* times the standard deviation of the slice *Z. α* =0.04 controls the strength of the noise relative to the natural intensity variation in that slice. Multiplying by *M*_*z*_ ensures the noise is added only to brain regions (mask = 1) and the background stays unchanged (mask = 0). The perturbed data *Ĩ* is used to fine-tune the pre-trained model defined as:

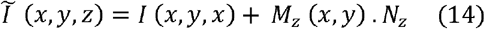

**Figure 5:**
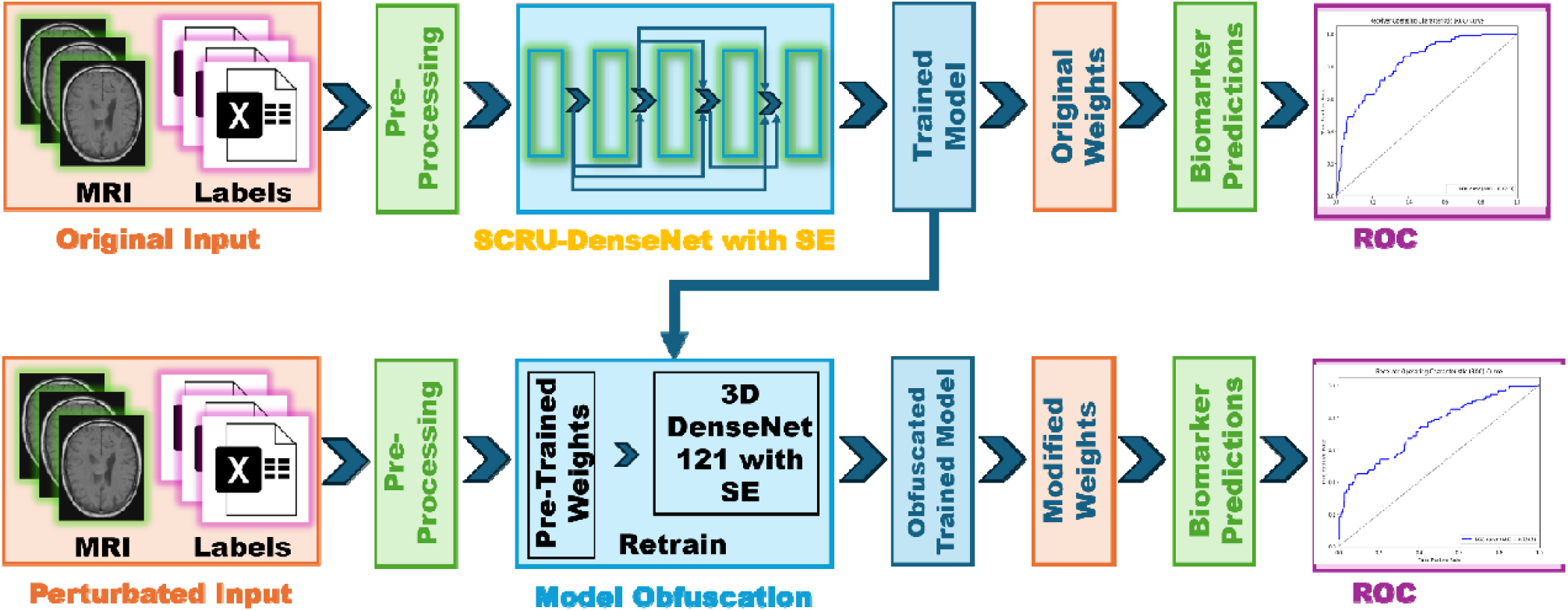
GlioVision model obfuscation

The resulting perturbed volume *Ĩ* is then used to fine-tune the original pre-trained network 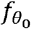 by solving:

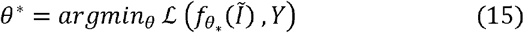

This additional fine-tuning step updates the model parameters from θ_0_ to θ_*_, guiding the network to shift its learned representations away from the exact distribution of the original training data while preserving features relevant to the diagnostic task. Here,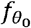 and 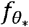 denote the models before and after fine-tuning, respectively.

The perturbed input data is represented as *Ĩ* while *I* denotes the original unperturbed data. The goal of this process is to ensure that the fine-tuned model 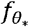 maintains high diagnostic accuracy, yet reduces the probability of accurately reconstructing the original sensitive images *X* from its internal representations. Formally, this privacy enhancement is expressed as a significant reduction in the conditional probability P defined as follows:

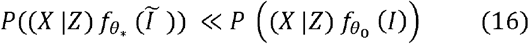

where *P*(*X* |*Z*) denotes the likelihood of reconstructing the original data *X* given the model’s internal representation *Z* where the model keeps diagnostic features but loses identifiable details. This inequality indicates that after fine-tuning on the perturbed data Ĩ, the model’s parameters contain substantially less identifiable information about the original patient images compared to the original model trained on *I*. To validate that privacy preservation does not compromise clinical performance, we evaluated the obfuscated model 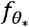 on both clean and perturbed test datasets for IDH mutation prediction across 92 patients, the results are presented in Figure 4 (b, c, d, e). For the majority class (class 0), the obfuscation induced only slight decreases in precision and F1 score, resulting in overall performance that remains relatively stable. The observed decrease for the minority class primarily reflects the natural class imbalance in IDH mutation status, where fewer positive samples naturally amplify variability in recall and F1 under obfuscation. Further experimental validation results are provided in the Supplementary document (sections S2, S3, and S8).

### 3.5. Ablation Study of Architectural Dependencies

This ablation study was designed to systematically measure performance in terms of IDH mutation accuracy, serving as the primary metric and contributing to the understanding of each new component added to our GlioVision architecture, represented in Figure 6. Starting with the standard 3D DenseNet baseline, we observed an initial accuracy of 0.687. The first major improvement was achieved by SCRU-DenseNet, which immediately elevated the accuracy to 0.809. This substantial jump validates the need for the SCRU module to efficiently extract and prioritize pathology-sensitive features from the volumetric input. Next, including the ACPS with its modality-specific training further refined the feature space, yielding an accuracy of 0.856, demonstrating the significant value of contextual integration. The largest reported gain was observed with the confidence-based CFPM, which increased the accuracy of the retained, high-confidence predictions to 0.9221. While this confirms the CFPM’s effectiveness in filtering out ambiguous cases to maximize diagnostic certainty, the final, total accuracy of the full GlioVision framework across the entire cohort stabilized at 0.8729. The SCRU-DenseNet’s parameter efficiency (approx. 12.2M parameters) on the most demanding tasks, demonstrating that GlioVision does not require high-end industrial clusters for deployment.

**Figure 6:**
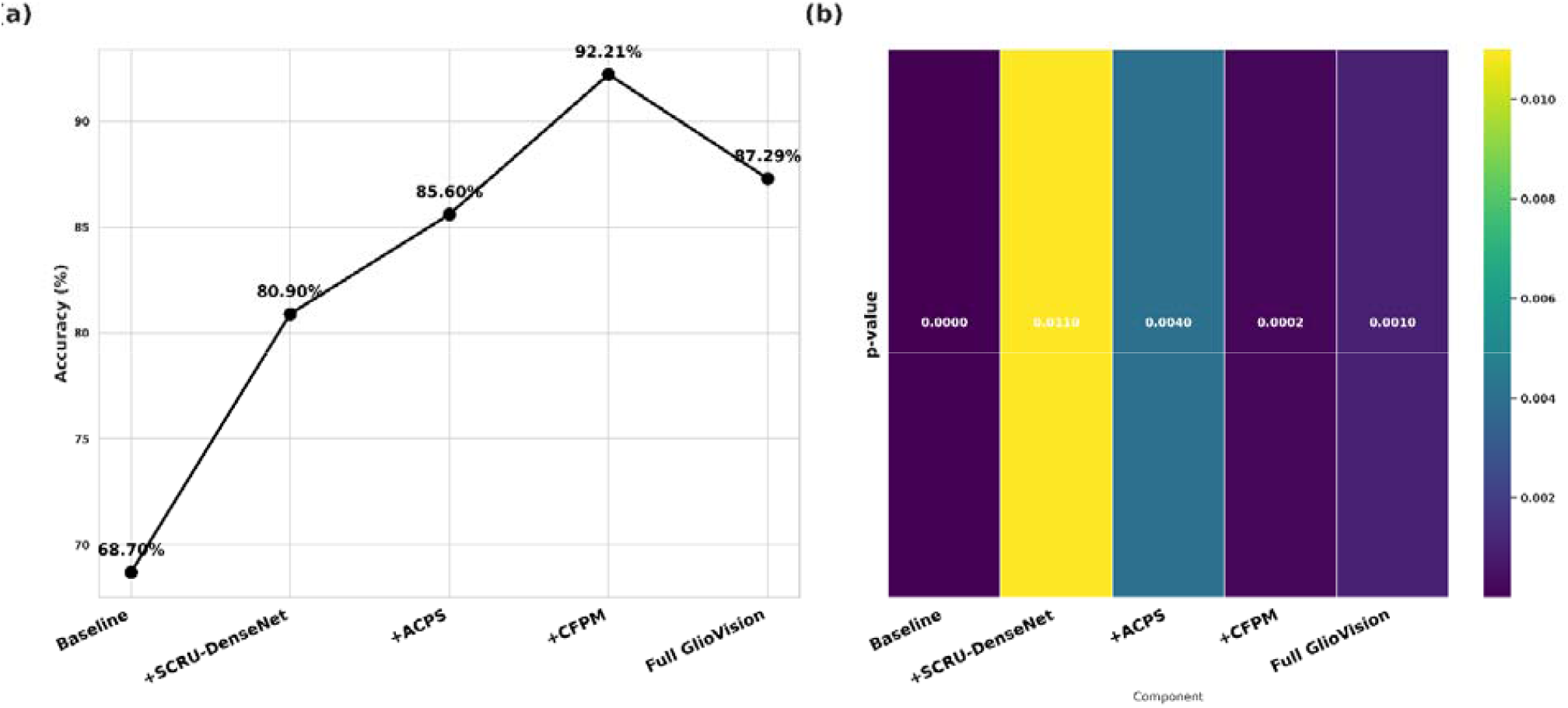
(a) Accuracy progression curve showing improvements from Baseline through +SCRU-DenseNet, +ACPS, +CFPM, to Full GlioVision, with exact accuracy values annotated. (b) A heatmap of p-values indicates the statistical significance of each component relative to the baseline, with lower values highlighting stronger significance.

## Discussion

Glioma survivors often experience reduced productivity and income in addition to the medical debt, and the society’s economy suffers from lost revenue due to productivity losses and premature deaths [29]. Finally, patients and their caregivers face reduced quality of life due to neurocognitive deficits and mental health impacts resulting from the tumors and their treatments. Therefore, improving glioma treatment is imperative to reduce the human and economic impacts of brain cancer. Our proposed method contributes to personalized care that improves outcomes and reduces unnecessary risks associated with cancer therapy and surgical intervention, and it has significant potential to alleviate the global healthcare burden. The model obfuscation approach helps retain the structural integrity and diagnostic relevance of the images, while preserving them largely intact and subtly perturbing features that might otherwise encode identifiable information into the model weights. At this stage, the network adapts its weights away from the exact distribution of the original training data, effectively blending the learned representations.

The GlioVision model is specifically designed to prioritize the prediction of key molecular biomarkers important for diagnosis and prognosis, including IDH and 1p/19q statuses, which support integrated tumor subtyping. Although the primary task of GlioVision is biomarker prediction, we still follow the WHO 2021 guidelines to interpret glioma subtyping based on the predicted molecular features. As shown in Table 4, If a case previously labeled as Grade 2 is predicted by the model as Grade 4 and simultaneously identified as IDH-wildtype, the prediction is consistent with the WHO 2021 definition of molecular glioblastoma, which prioritizes molecular features over histology. The translational relevance of GlioVision is further shown by its performance in the clinical cohort with known molecular features. By identifying R132H/R132C mutations at the genomic level, we confirmed that the model’s predictions align with the WHO 2021 integrated diagnosis.

Our dataset was compiled from multiple institutions using publicly available cohorts, including The Cancer Imaging Archive (TCIA), as mentioned in subsection 2.4 above. For each biomarker (IDH, 1p/19q, MGMT, and tumor grade), the test set was strictly held out during model development and consists of samples originating from multiple institutions, ensuring that the evaluation reflects heterogeneous clinical imaging data rather than a single-center distribution, as summarized in Supplementary document Table 3. In addition to this multi-institutional evaluation, we further explored a separate clinical cohort for supplementary assessment. This validation strategy, combining a strictly unseen multi-institutional test set with an additional independent clinical dataset, helps mitigate the risk of performance inflation due to institutional bias and demonstrates that the model’s performance is not limited to a single data source.

Our tests on the feasibility of privacy protections for MRI images revealed a major challenge in protecting training datasets while also protecting intellectual property rights for the researchers and developers. We anticipate that these issues will influence data-sharing considerations for imaging datasets in the near future. To accommodate these issues, we have maintained two versions of the proposed algorithm: one that includes obfuscated GlioVision and one that does not, which retains the full power of the training datasets. The proposed MRI modality-specific training technique exploits the diverse, complementary information intrinsic to multi-parametric MRI to improve generalization performance in the presence of tumor variability and scanner differences. To address domain shifts arising from cross-database variation, including differences in acquisition protocols, scanner hardware, and noise characteristics, we integrate the SE block after the final dense block, thereby minimizing computational overhead while leveraging fully aggregated features. The embedding is jointly learned with the network to reflect domain-specific variations. This fusion mechanism enables recalibration of weights to adaptively adjust domain-specific factors, facilitating the model’s selective scaling of channel features in accordance with dataset variability.

The GlioVision framework has natural extensions that will be addressed in future work. A major goal is to derive tumor volumes (contrast-enhancing on T1-GD and hyperintense on FLAIR), which serve as clinically meaningful biomarkers. This can open the future foundation for downstream tasks such as predicting overall survival and treatment response (e.g., to radiotherapy/XRT or chemotherapy). While these outcome models are not included in the present manuscript, they could be a reasonable extension for future work, particularly with access to public glioma data cohorts that are aligned with the 2021 WHO CNS classification [2].

GlioVision presents an automated pipeline for non-invasive glioma characterization, uniting biomarker prediction and privacy-preserving features within a confidence-controlled reliability module. Trained on the most extensive multi-cohort glioma dataset available, GlioVision achieves improved performance across IDH mutation, 1p/19q co-deletion, MGMT methylation, and WHO grade prediction, while maintaining high sensitivity, specificity, and reliability, as validated on independent clinical cohorts. By providing modality-specific training strategies, pre-trained weights, a codebase model, and a resource-efficient solution, GlioVision establishes a practical foundation for future applications beyond those focused on glioma, enabling real-time clinical use. Unlike traditional radiomics, which utilize fixed mathematical formulas to extract texture and shape features, the deep features generated by the SCRU-DenseNet are learned directly from the multi-parametric MRI manifold. This approach offers superior biological relevance by capturing hierarchical patterns of tumor heterogeneity.

## Conclusion

GlioVision’s deep learning framework presents a pipeline for non-invasive characterization of key molecular biomarkers and tumor grade, uniting biomarker prediction and privacy-preserving features within a confidence-controlled reliability module. Trained on the extensive multi-cohort glioma dataset, GlioVision achieves strong performance across IDH mutation, 1p/19q co-deletion, MGMT methylation, and WHO grade prediction, with high sensitivity, specificity, and reliability, as validated on independent clinical cohorts. Our study focuses on accurate molecular biomarker prediction while aligning with current WHO guidelines, providing a reliable foundation for future integrated glioma subtyping. By providing a modality-specific training strategy, an accessible codebase model, and a resource-efficient design, GlioVision introduces a practical foundation for future applications beyond glioma-focused tasks.

## DATA AND CODE AVAILABILITY

The datasets analyzed in the current study are available in the Cancer Imaging Archive (TCIA) and the Erasmus Glioma Database (EGD). These datasets are publicly available to researchers upon standard user registration and data access request via the respective repository platforms. The datasets can be accessed via the following persistent web links:

UCSF-PDGM: https://www.cancerimagingarchive.net/collection/ucsf-pdgm/

UPENN-GBM: https://www.cancerimagingarchive.net/collection/upenn-gbm/

TCGA-LGG: https://www.cancerimagingarchive.net/collection/tcga-lgg/

TCGA-GBM: https://www.cancerimagingarchive.net/collection/tcga-gbm/

IvyGAP: https://www.cancerimagingarchive.net/collection/ivygap/

RHUH-GBM: https://www.cancerimagingarchive.net/collection/rhuh-gbm/

EGD: https://xnat.health-ri.nl/

The source code of this study is provided as a ZIP archive in the Supplementary/Related files section with this manuscript.

## CRediT statement

**AN** is involved in the overall study design, model development, coding implementation, and experimental analysis. **MNC** contributed to data processing and assisted in the technical execution and result validation. AH **s**erved as the principal supervisor, providing overall guidance, conceptualization of the study, and methodological direction. **ASH** provided clinical label verification, which is essential for model evaluation. **JZ** served as the clinical collaborator, validating the study’s medical relevance and verifying clinical. **XJ** provided computational resources and critical feedback during manuscript review. **YH** facilitated access to clinical data through the affiliated hospital.

