## Supplementray document for "GlioVision: A Multi-Modal MRI Framework for Non-Invasive Glioma Molecular Biomarkers Prediction"

**(Supplementary Document)**

Anam Nazir^*^, Muhammad Nadeem Cheema, Yu-Chun Hsu, Xiaoqian Jiang, Jay-Jiguang Zhu, Akdes Serin Harmanci, Arif Harmanci

Anam Nazir, Muhammad Nadeem Cheema, Yu-Chun Hsu, Xiaoqian Jiang, and Arif Harmanci are with the Department of Health Data Science and Artificial Intelligence, D. Bradley McWilliams School of Biomedical Informatics, The University of Texas Health Science Center, Houston, TX, 77030, USA. (Corresponding)

Jay-Jiguang Zhu is with the Department of Neurosurgery, The University of Texas Health Science Center, Houston, TX, 77030, USA.

Akdes Serin Harmanci is with the Department of Neurosurgery, Baylor College of Medicine, Houston,TX 77030, USA

**S1. Adaptive Contrast-specific Processing Stream (ACPS) and SCRU-DenseNet Architectural Framework**

The ACPS SCRU-DenseNet algorithm, as implemented in the GlioVision pipeline, facilitates the automated molecular stratification of gliomas by integrating multi-parametric MRI volumes through a structured feature-fusion approach. The workflow initiates with the Adaptive Contrast-specific Processing Stream (ACPS) path, where each imaging modality (T1, T1Gd, T2, and FLAIR) is concatenated with its corresponding anatomical mask and processed via independent encoders to extract modality-specific features. These features are then integrated into a global feature map through a fusion step, followed by a *1x1x1* convolutional harmonization layer that aligns the high-dimensional data for the primary processing stage. The SCRU-DenseNet backbone employs dense connectivity and residual skip connections to mitigate the vanishing gradient problem and ensure efficient feature propagation throughout the network. This architecture ultimately computes a multi-task output, generating the clinical probability distributions *P* for four critical biomarkers: IDH mutation, 1p/19q codeletion, MGMT methylation, and WHO Grade. The following pseudocode explains the implementation of this module in the proposed GlioVision model:

**Algorithm S1:**

*1****. Input:*** *Multi-parametric volume V = {x_i_, m} where x is the sequence and m is the mask.*

*2.* ***ACPS Path:*** *Фi = Encoder_i_ (Concatenate {x_i_, m}) for each modality i.*

*3.* ***Fusion****: Ф= Concatenate (Ф_T1_, Ф_T1Gd_, Ф_T2_, Ф_FLAIR_).*

*4.* ***Harmonization:*** *Ф^= Conv _1X1X1_ (Ф).*

*5.* ***Backbone:*** *Y = {SCRU-DenseNet (Ф^) with residual skip connections.*

*6.* ***Output:*** *Biomarker probabilities P(IDH), P(1p/19q), P(MGMT), P(Grade).*

**S2. Statistical Verification of Model Privacy**

To quantify the privacy-utility trade-off, we analyzed the model's prediction confidence (C) across varying levels of Gaussian noise (2% and 5%). In our experiment depicted in Figures 1a, b, and 2, along with Table 1, injecting 5 % Gaussian noise into MR images led to a statistically significant decrease in average prediction confidence, while accuracy remained effectively unchanged under in-distribution conditions. Specifically, independent‑sample t-tests comparing $C_{clean}$ and $C_{noisy}$ (5 %) yield t = 6.4129 on the training cohort and t = 4.7756 on the holdout cohort, confirming a highly reliable confidence. Notably, the comparison between 2 % and 5 % Gaussian noise also shows significant effects, with t = 4.5845 for the train and t = 3.4866 for the holdout, indicating that the confidence gap widens with increasing noise magnitude. Conversely, $C_{clean}$ vs $C_{noisy}$ at 2 % fails to reach significance in holdout (t = 0.9049), implying that light augmentation preserves model certainty. These findings validate our privacy-preserving watermarking hypothesis: controlled Gaussian noise yields a measurable reduction in confidence, yet the model continues to accurately predict a desirable property for detecting potential data leakage without impairing predictive performance.

Table 1: Statistical Comparison of Prediction Confidence under Noise Perturbation

| Cohort | Model Comparison | t-value | p-value | Significance |
| --- | --- | --- | --- | --- |
| Train | Clean vs 2% Noise | 1.7219 | 0.08552 | No |
| Train | Clean vs 5% Noise | 6.4129 | 2.601e-10 | Yes |
| Train | 2% Noise vs 5% Noise | 4.5845 | 5.375e-06 | Yes |
| Holdout | Clean vs 2% Noise | 0.9049 | 0.3689 | No |
| Holdout | Clean vs 5% Noise | 4.7756 | 1.083e-05 | Yes |
| Holdout | 2% Noise vs 5% Noise | 3.4866 | 0.0008895 | Yes |

**S3. Privacy Safeguards for assessing re-identification risk**

To assess the re-identification risk of our privacy mechanism, we further investigated whether advanced augmentation techniques could intentionally mask this confidence shift, as shown in Figure 3. First, we applied adversarial perturbations using the Fast Gradient Sign Method (FGSM) [1]. This did not effectively conceal data reuse. The gap *ΔC*${=C}_{clean}-C_{noisy}$ remained, and classification accuracy declined under adversarial noise. Next, we tested Mixup and CutMix [2] augmentations, which synthesize new MRIs by interpolating or splicing regions from multiple scans. These blending strategies partially narrowed the confidence gap but produced visible artifacts, limiting their ability to fully hide data provenance.

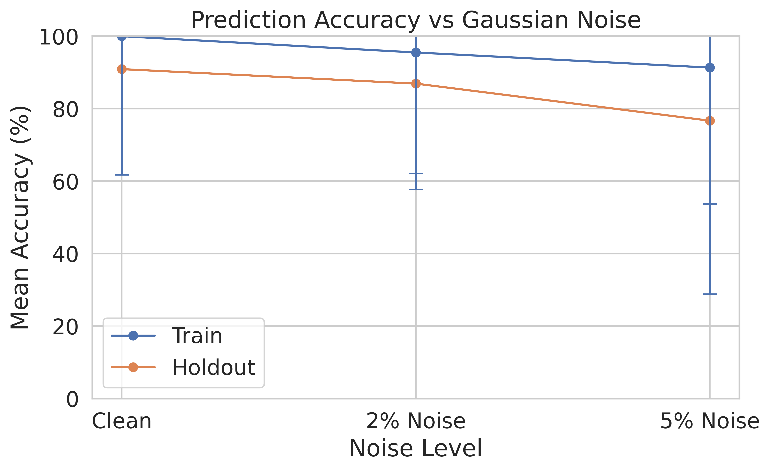

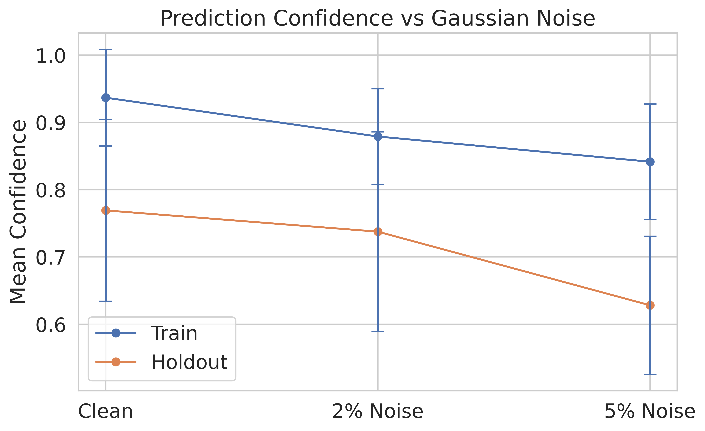

Figure 1: a) Confidence score vs Gaussian noise, b) Prediction accuracy vs Gaussian noise

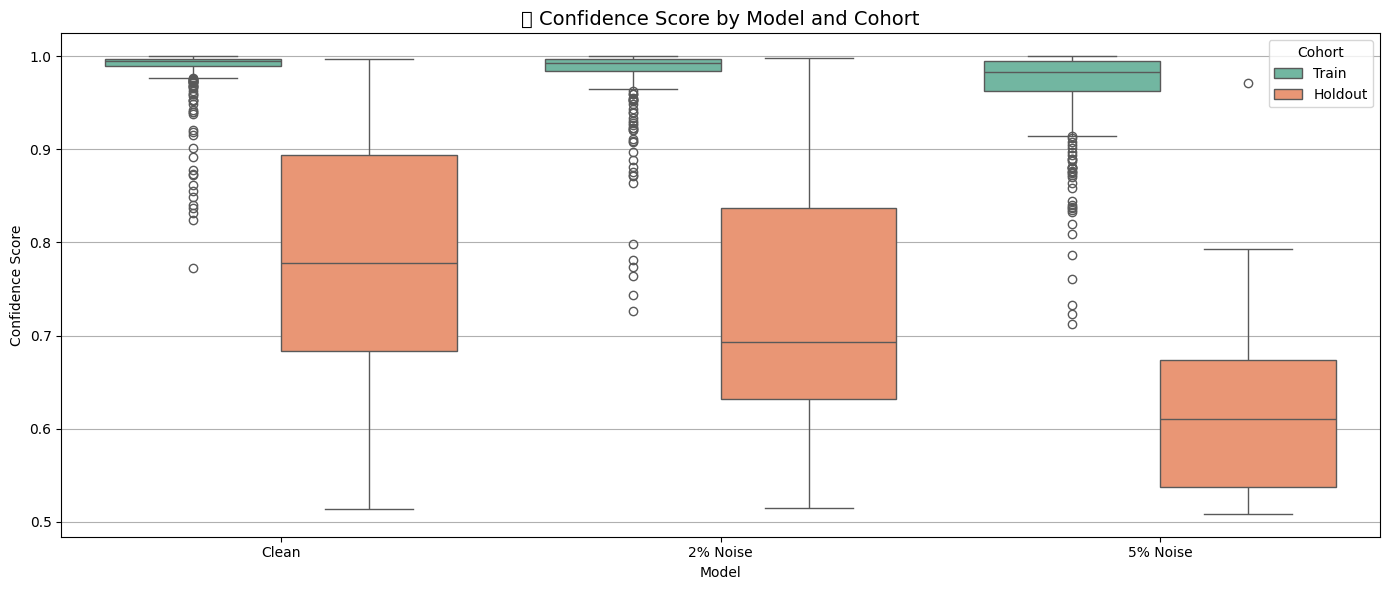

Figure 2: Train and hold out data, confidence, and statistical visualization

In this preliminary experiment, the most effective camouflage was achieved by adding Gaussian noise, shown in Figure 3, directly to the training images at a controlled level of (2-5%) together. This simple perturbation preserves anatomical structure while deliberately modifying pixel-level distributions. As a result, the trained model shows nearly the same confidence levels for both training and holdout data under noise: $C_{noisy} \approx C_{clean}$. Hence, we formulated a hypothesis based on this finding: Controlled Gaussian noise augmentation applied during training (e.g., at 2–5% intensity) enables a trained model to exhibit similar confidence distributions on clean and noisy data, effectively camouflaging the confidence shift (ΔC ≈ 0) while preserving classification accuracy. This hypothesis remains provisional and requires further experiments for formal validation. This phase of the study frames an initial, tentative hypothesis to guide further experimentation and statistical validation. By explicitly defining the independent variable (controlled Gaussian noise at 2–5 % intensity) and the dependent outcomes (confidence shift ΔC

 
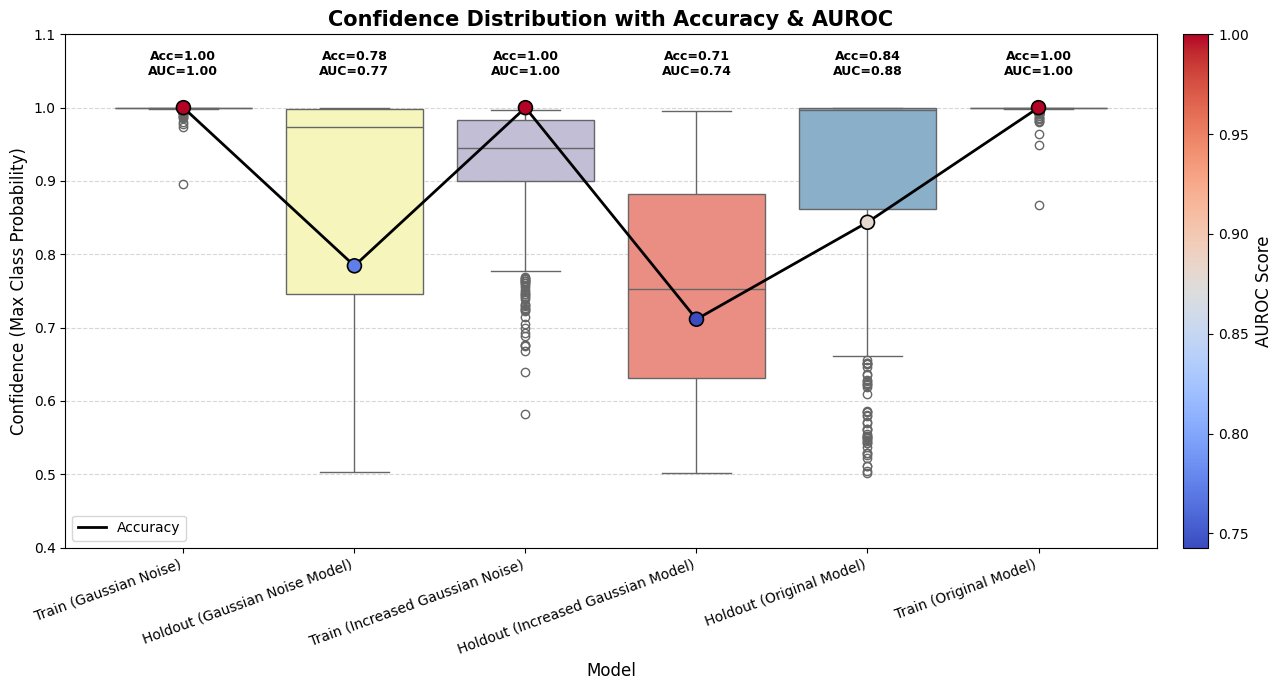

Figure 3: Privacy camouflage experiments

and classification accuracy), this working hypothesis establishes clear parameters for replication. Together, these findings highlight a critical trade-off. Our controlled noise-based confidence test provides a simple, non-intrusive safeguard for verifying the independence of training data in collaborative glioma biomarker research. However, the same noise augmentation, if misapplied, can serve as a privacy camouflage, concealing test-set misuse.

**S4. Cross-Institutional Data Handling**

The 70/15/15 subject-level split was purposefully applied to a pooled multi-institutional dataset to ensure that the ACPS architecture was exposed to a diverse range of magnetic field strengths (1.5T and 3T) and vendor-specific acquisition protocols during the learning phase. By incorporating subjects from six distinct international cohorts (UCSF-PDGM [3], UPENN-GBM [4], TCGA-LGG [5], TCGA-GBM [6], EGD [7], IvyGAP [8], and RHUH-GBM [9]) into the training part of the dataset, the model was forced to converge on institutional-invariant radiogenomic signatures rather than memorizing site-specific artifacts. The resulting internal test sets comprising 278 subjects for IDH and 127 for 1p/19q in Tables 2 and 3, respectively, therefore represent a heterogeneous mini-set of unseen data. This validation, coupled with a strictly independent external holdout from a seventh clinical data cohort, mitigates the risk of performance inflation from institutional bias.

Table 2: Data partitioning strategy with Institutional Diversity

| Data Cohort | Subjects (N) | IDH (0/1) | MGMT (0/1) | 1p/19q (0/1) | Grade (2/3/4) |
| --- | --- | --- | --- | --- | --- |
| EGD | 774 | 312 / 155 | 0 / 0 | 186 / 73 | 135 / 79 / 502 |
| TCGA (LGG/GBM) | 164 | 90 / 56 | 0 / 0 | 147 / 12 | 27 / 37 / 100 |
| UCSF-PDGM | 491 | 390 / 101 | 296 / 112 | 385 / 15 | 55 / 42 / 394 |
| UPENN-GBM | 671 | 546 / 125 | 121 / 170 | 0 / 0 | 0 / 0 / 0 |
| RHUH-GBM | 40 | 36 / 4 | 0 / 0 | 0 / 0 | 0 / 0 / 40 |
| IvyGAP | 39 | 31 / 6 | 0 / 0 | 27 / 3 | 0 / 2 / 36 |
| TOTAL | 2,279 | 1,405/447 | 417/282 | 745/103 | 254/180/1,103 |

Table 3: Dataset Subject-level Split for Four Biomarkers

| Biomarker | Total Available (N) | Training (70%) | Validation (15%) | Test (15%) |
| --- | --- | --- | --- | --- |
| IDH Status | 1,852 | 1,296 | 278 | 278 |
| 1p/19q Status | 848 | 594 | 127 | 127 |
| MGMT Status | 699 | 489 | 105 | 105 |
| Tumor Grade | 1,537 | 1,076 | 230 | 231 |

**S5. Automated Tumor Mask Extraction**

To ensure spatial consistency across heterogeneous data sources, we developed an automated preprocessing pipeline for extracting Whole Tumor (WT) masks. We used a specialized 3D U-Net architecture [10], pre-trained on the BraTS (Brain Tumor Segmentation) [11] challenge dataset, for voxel-wise segmentation. The U-Net follows an encoder-decoder structure with skip connections, enabling the capture of both high-resolution local features and global contextual information. For each subject, the model processed the multi-parametric MRI stack (T1, T1Gd, T2, and FLAIR) to generate a unified WT mask, defined as the union of the contrast-enhancing core, non-enhancing/necrotic regions, and peritumoral edema. This automated approach was applied consistently to all 2,279 public subjects and the 57 local clinical cases, followed by a manual verification step to ensure anatomical accuracy before the masks were fed into the GlioVision predictive manifold.

**S6. CFPM Performance Impact and Omission Statistics**

The systematic application of CFPM across all biomarker tasks confirms the framework’s reliability. For 1p/19q codeletion and MGMT methylation, the mechanisms omitted 16.5% and 17.1 % of cases, respectively. These omitted cases were characterized by high entropy in the posterior probability distribution, often corresponding to atypical imaging presentations. By excluding these ambiguous outputs, the accuracy of MGMT prediction increased from 87.09% to 93.10% as shown in Table 4. Crucially, the omission rate did not exceed 20% across any task, ensuring the model remains practically useful in a high-volume clinical workflow while significantly reducing the risk of automated misdiagnosis.

Table 4: Statistical Comparison of Prediction Confidence under Noise Perturbation

| **Prediction Task** | **Total Samples (N)** | **Omitted Cases (N)** | **Omission Rate (%)** | **Accuracy (Before CFPM)** | **Accuracy (After CFPM)** |
| --- | --- | --- | --- | --- | --- |
| IDH Mutation | 92 | 15 | 16.3% | 85.87% | 92.21% |
| 1p/19q Codeletion | 127 | 21 | 16.5% | 83.33% | 89.62% |
| MGMT Methylation | 105 | 18 | 17.1% | 87.09% | 93.10% |
| Tumor Grade | 150 | 28 | 18.6% | 87.83% | 94.26% |

**S7. Clinical Deployment and Workflow Integration:**

To bridge the gap between retrospective evaluation and clinical utility, GlioVision incorporates a fail-safe deployment protocol. In this workflow, the model serves as an automated triage tool; high-confidence predictions (as determined by the CFPM) [12] provide immediate decision support for surgical planning, while 'omitted' cases are flagged for mandatory confirmatory genomic testing and expert neuroradiological review. This dual-track system addresses the challenge of clinical uncertainty and prevents the misuse of low-certainty outputs. Furthermore, the integration of DTI-A ensures that this deployment remains compliant with institutional privacy standards, allowing for secure multi-center validation without exposing sensitive patient data. By treating AI as a collaborative 'second reader' that knows its own limitations, GlioVision establishes a realistic pathway for integration into modern neuro-oncological diagnostic pipelines as shown in Figure 4.

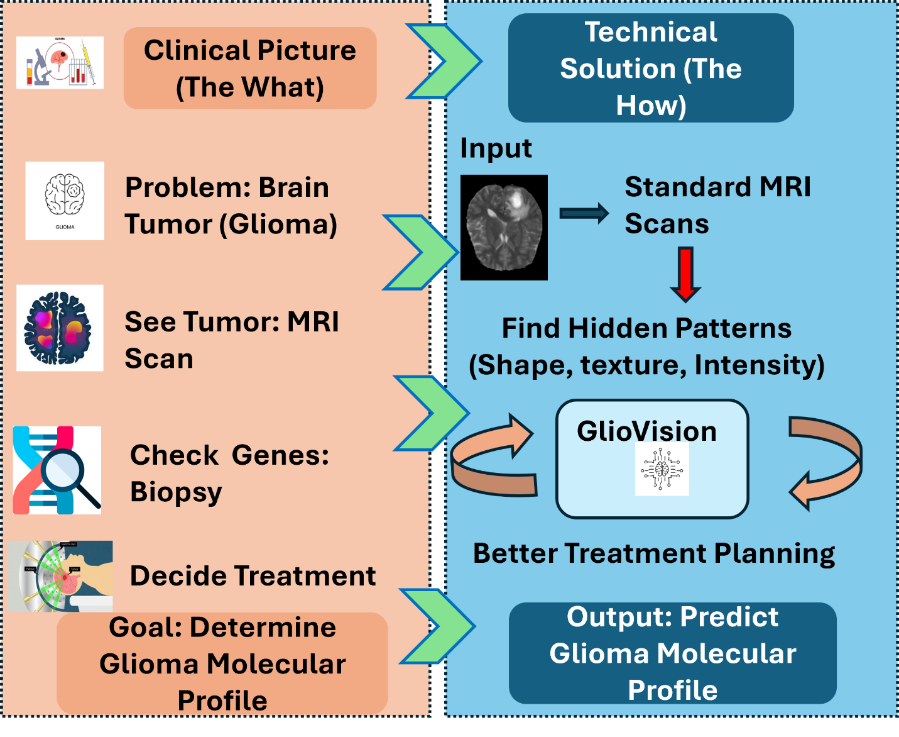

Figure 4: Clinical Deployment and Workflow Integration of GlioVision

**S8. Validation of DTI-A Privacy Strategy**

The selection of controlled Gaussian noise coupled with a fine-tuning (transfer learning) regime is a deliberate architectural choice designed to balance clinical utility with biometric de-identification. Standard de-identification often fails because neural networks can "memorize" high-frequency structural textures (cortical folding, vascular patterns) that act as biometric identifiers. Unlike conventional privacy approaches, which focus primarily on secure storage or differential privacy techniques, our strategy directly reshapes the model’s learned patterns by retraining on carefully perturbed samples, reducing the risk of proprietary or patient-identifiable image data being inferred from the model’s weights.

First, the baseline model is trained on the original, high-quality multi-parametric MRI input dataset to learn representative features for prediction tasks. After the initial model has been trained, its weights serve as the foundation for obfuscation. Subsequently, a perturbed version of the dataset is generated by adding carefully controlled Gaussian noise exclusively to brain tissue regions, without altering the background, thereby preserving global anatomical structures. To define the model obfuscation process, we have generated a modified version of the dataset by adding controlled Gaussian noise exclusively to brain tissue regions, leaving the background unchanged to preserve global anatomical context. To ensure this privacy layer is clinically viable, we performed a comparative analysis across clean and perturbed test sets.

Table 5 : Statistical Comparison of Prediction Confidence under Noise Perturbation

| **Metric (IDH Class 0)** | **Original Model (θ_0_)** | **Obfuscated Model (θ_∗_)** | **Delta (Δ)** |
| --- | --- | --- | --- |
| **Precision** | 0.88 | 0.86 | -0.02 |
| **F1-Score** | 0.91 | 0.89 | 0.02 |
| **Specificity** | 0.78 | 0.77 | -0.01 |

The results in Table 5 confirm that the obfuscation induced only marginal fluctuations in performance. The slight variability in the minority class recall is a known effect of class imbalance, yet the overall diagnostic accuracy remains within the threshold required for clinical decision support.

To confirm that the DTI-A process induces a meaningful change in the model's internal representation, we analyzed the weight distributions of the final convolutional layers. Figure 5 displays the comparative histograms of the model weights before and after fine-tuning. The observable shift and redistribution of parameter values confirm that the network has adapted to the perturbed input manifold. This adaptation reduces the likelihood of data reconstruction by shifting the parameters from the clean model to the obfuscated model. The model no longer has a direct mathematical mapping to the original patient’s unique anatomical 'fingerprint,' effectively de-identifying the training data at the architectural level.

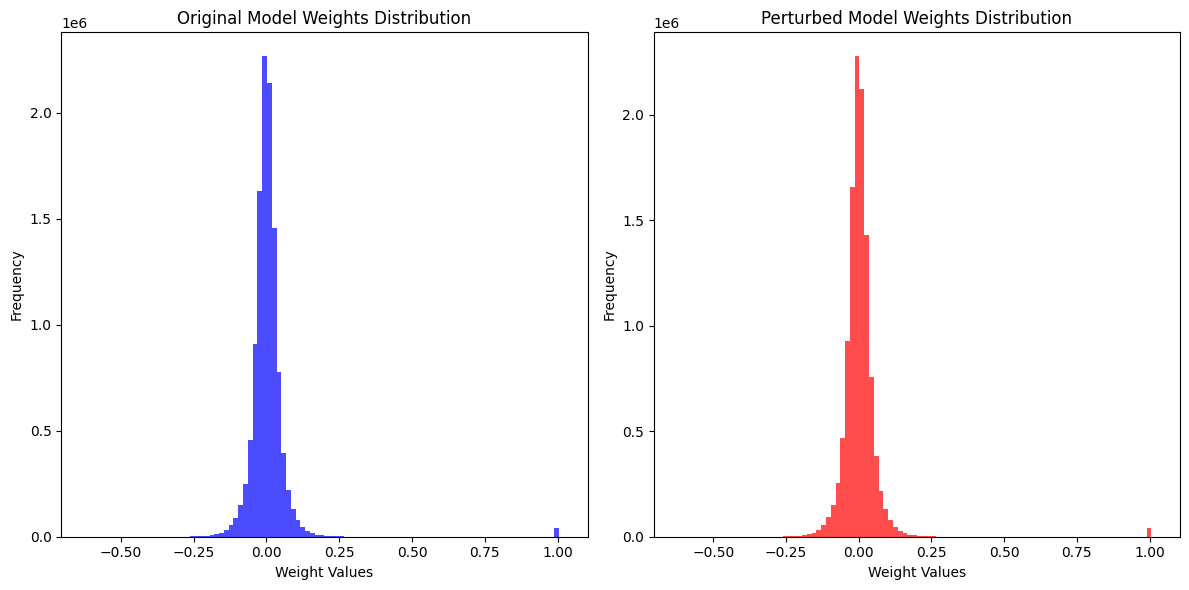

Figure 5: Clean GlioVision and obfuscated GlioVision model weights for showing the difference after data perturbations
